# Stable characteristics of intrapopulation heterogeneity in virus-specific Th1 cells during chronic viral challenge infection

**DOI:** 10.1101/2025.09.15.676263

**Authors:** Valerie Plajer, Adrián Madrigal-Avilés, Maria Dzamukova, Nayar Durán-Hernández, Philippe Saikali, Vivien Holecska, Isabel Panse, Katrin Lehmann, Jinfang Zhu, Mir-Farzin Mashreghi, Ahmed N. Hegazy, Caroline Peine, Max Löhning

**Affiliations:** Charité – Universitätsmedizin Berlin, corporate member of Freie Universität Berlin and Humboldt-Universität zu Berlin, Department of Rheumatology and Clinical Immunology, Experimental Immunology and Osteoarthritis Research, 10117 Berlin, Germany; Pitzer Laboratory of Osteoarthritis Research, German Rheumatology Research Center (DRFZ), A Leibniz Institute, 10117 Berlin, Germany; Josep Carreras Leukaemia Research Institute (IJC), 08916 Badalona, Barcelona, Catalonia, Spain; Therapeutic Gene Regulation, German Rheumatology Research Center (DRFZ), A Leibniz Institute, 10117 Berlin, Germany; Laboratory of Immune System Biology, National Institute of Allergy and Infectious Diseases, National Institutes of Health, Bethesda, MD 20892, USA; German Center for Child and Adolescent Health (DZKJ), 10117 Berlin, Germany; Charité – Universitätsmedizin Berlin, corporate member of Freie Universität Berlin and Humboldt-Universität zu Berlin, Department of Gastroenterology, Infectious Diseases and Rheumatology, 12203 Berlin, Germany; Laboratory of Inflammatory Mechanisms, German Rheumatology Research Center (DRFZ), A Leibniz Institute, 10117 Berlin, Germany

**Keywords:** CD4 T cells, Th1 cells, T-bet, quantitative stability, chronic infection, exhaustion

## Abstract

Virus-specific CD4^+^ T cells typically undergo T helper (Th) 1 differentiation and contribute to a type 1 immune response in infection with lymphocytic choriomeningitis virus (LCMV). Using this model pathogen, we performed an in-depth analysis of the quantitative expression stability of the Th1 key transcription factor T-bet. Previously, it was shown that virus-specific Th1 cells arising in acute infections expressed T-bet at distinct intensities and maintained their T-bet expression differences after viral clearance as memory cells for weeks in the steady state. However, it was unclear whether differential T-bet expression was associated with heterogeneity inside the Th1 population and if the quantitative T-bet memory, particularly of those cells expressing T-bet at low levels, could withhold the strong and continuous stimulation present during chronic infection.

Using T-bet-ZsGreen reporter mice, virus-specific Th1 cells were characterized phenotypically at protein, RNA, and DNA/chromatin accessibility levels. The Th1 cells arising during acute LCMV Armstrong infection showed a continuous spectrum of T-bet expression, ranging from cells with very high T-bet to cells with low T-bet. Even though the cells with low T-bet expression clearly possessed Th1 characteristics, they additionally showed certain Tfh-like features at protein and RNA level. When virus-specific Th1 cells were sorted according to T-bet-ZsGreen reporter expression intensity, adoptively transferred, and rechallenged by infecting the host animals with the chronic Clone 13 strain of LCMV, they maintained quantitative differences in T-bet reporter and IFN-γ expression levels. A subpopulation of the progeny of the former T-bet^low^ cells still showed a mild Tfh-associated phenotype. Independent of their past and present T-bet expression level, all virus-reactive CD4^+^ T cells acquired phenotypic signs of exhaustion as characterized by upregulation of PD-1, LAG3, and TOX and vast absence of effector cytokine co-expression in the chronic infection environment.

Collectively, our findings highlight the heterogeneity of T-bet^+^ antiviral CD4^+^ T cells and the stability of quantitative differences in individual virus-specific CD4^+^ T cells during chronic viral challenge infection.

## 1 Introduction

When CD4^+^ T cells become activated, they can differentiate into particular subsets depending on environmental cues. During intracellular infections, as it is the case for viruses, the reactive cells often acquire features of the Th1 lineage, which includes the expression of the master-regulator transcription factor T-bet and the signature effector cytokine interferon-gamma (IFN-γ). Previously, we have shown that quantitative differences in the expression levels of T-bet and IFN-γ can be stably maintained in memory Th1 cells at steady state. Furthermore, in differentiated Th1 cells, T-bet amounts quantitatively regulate IFN-γ production (1).

Apart from Th1 cells, also T follicular helper (Tfh) cells can arise during viral infections. The expression of BCL-6, their key transcription factor, induces the upregulation of CXCR5 and the consequent homing to the B cell follicle (2). There, they can drive the formation of germinal centers (GC) and support isotype class switching of B cells (3). For Tfh cells, the relevance of T-bet and IFN-γ expression has been previously studied. Transient T-bet expression has been shown to take place during GC Tfh differentiation, which allows these cells to maintain an accessible IFN-γ locus and produce this cytokine even in the absence of T-bet (4). This promotes the isotype class switch of B cells towards IgG2a/IgG2c, which is beneficial for viral clearance (5–7).

During chronic infections, T cells undergo functional adaptation that may include exhaustion whereby they make multiple phenotypic changes to adjust to the persistent antigen exposure and the consequent inflammatory environment (8). Lymphocytic choriomeningitis virus (LCMV) is a commonly used virus model to study both acute and chronic infections side-by-side in mice using the different substrains, LCMV Armstrong and Clone 13, respectively (9). As CD4^+^ T cell help is important to promote effector functions of CD8^+^ T cells and B cells during chronic infections, it has been previously assessed whether a vaccine-induced CD4^+^ T cell response could have a protective effect in chronic viral infections (10). However, during chronic LCMV infections severe immunopathology was reported caused by the vaccine-primed cells, which were recovered in high numbers from multiple organs, maintained a Th1 phenotype, and exhibited reduced features of exhaustion compared to CD4^+^ T cells from unvaccinated animals. As this study focused on deciphering the cause for immunopathology, it remained unclear how heterogenous the vaccine-primed CD4^+^ T cells are and if their T-bet expression levels could influence their exhaustion potential. In hepatitis B and C infections, which can cause either acute or chronic infections in contrast to the universally chronic pathogens HIV or LCMV Clone 13, it has been described that high T-bet levels are observed in CD8^+^ T cells of spontaneously resolving, but not in chronically evolving infections (11). This suggests that high T-bet levels and the associated elevated expression of IFN-γ by at least CD8^+^ T cells can play a pivotal role in the outcome of the infection. Taken together and set in the context of vaccine development to prevent chronic infections, it is of interest to define if the T-bet levels of previously primed, virus-reactive CD4^+^ T cells could influence the exhaustion potential and thereby the functionality of these cells during a challenge with a viral strain inducing chronic infection.

We have previously demonstrated that T-bet expression levels induced during LCMV infection regulate the magnitude of IFN-γ expression and govern the plasticity of Th1 cells towards the Th2 lineage. We observed that LCMV infection induces heterogeneous T-bet expression in LCMV-specific CD4^+^ T cells. Furthermore, we showed that the observed T-bet expression gradient remains stable even after secondary acute LCMV challenge. Additionally, we have shown that the magnitude of T-bet expression regulates the plasticity of Th1 cells towards the Th2 phenotype, and this T-bet-dependent plasticity remains unaltered even after secondary infection (12).

Here, we used the murine T-bet ZsGreen reporter model to investigate the composition of the T-bet positive CD4^+^ T cell pool after an acute infection with LCMV. We found that although both T-bet^high^ and T-bet^low^ cells had acquired a Th1 phenotype, some of the T-bet^low^ cells showed additional characteristics associated with Tfh cells, suggesting an intrapopulation heterogeneity of virus-specific CD4^+^ T cells linked to T-bet expression levels. During chronic LCMV infection, quantitative differences in T-bet reporter and IFN-γ expression were maintained, but none of the cell types were protected from acquiring features of exhaustion. Our findings further highlight the stability of quantitative T-bet differences in CD4^+^ T cells and suggest that even high T-bet expression cannot prevent the acquisition of some exhaustion-associated features.

## 2 Materials & Methods

### 2.1 Mice

T-bet-ZsGreen reporter mice (13) were backcrossed to C57BL/6J background. Smarta1-TCR transgenic mice expressing a TCR specific for the LCMV glycoprotein (GP) 61-80 epitope (Smarta) (14) as well as Thy1.1 as a congenic marker were crossed to T-bet ZsGreen reporter mice and used as organ donors for the isolation of LCMV-specific CD4^+^ T cells. T-bet ZsGreen reporter mice with the congenic marker Thy1.2^+^ were used as recipients in adoptive cell transfer experiments. *In vivo* experiments were performed with male and female mice at the age of 8-20 weeks. Mice were bred under specific-pathogen free (SPF) conditions at the Charité animal facility, Berlin. Animal protocols were performed in accordance with the German law for animal protection and with permission from the local veterinary offices. All experiments were approved by the Landesamt für Gesundheit und Soziales in Berlin (LAGeSo, approval number G0205/18).

### 2.2 Adoptive T cell transfer and virus propagation, infection, and viral titer determination

Naive CD4^+^ T cells from T-bet ZsGreen Smarta Thy1.1^+^ mice were purified by magnetic cell sorting in a negative enrichment approach with biotin-labeled antibodies against CD8 (53-6.7), NK1.1 (PK136), CD11b (M1/70), CD11c (HL3), CD25 (7D4), Gr-1 (RB6-8C5), CD19 (1D3), and CXCR3 (CXCR3-173) in combination with anti-biotin microbeads according to the manufacturer instructions (Miltenyi Biotec). For primary infections, 2 × 10^5^ purified naïve Smarta Thy1.1^+^ CD4^+^ T cells were transferred i.v. into recipients one to five days before i.v. infection with 200 PFU (low dose) LCMV Armstrong (Arm). On day 10 post infection with LCMV Arm, the transferred cells were isolated from spleen and lymph nodes by mechanical disruption and either analyzed or positively enriched with magnetic anti-CD90.1 (Thy1.1) microbeads according to manufacturer’s instructions (Miltenyi Biotec). Subsequently, the cells were pooled from four to five mice and FACS sorted (untouched) according to their T-bet ZsGreen brightness into High and Low expressors as well as T-bet ZsGreen Mock-sorted live cells. These cells were either used for RNA (5×10^5^ cells/sample) and ATAC (1×10^5^ cells/sample) sequencing or used for challenge experiments. For challenge infections, 1-2 × 10^5^ T-bet ZsGreen-sorted CD4^+^ T cells were re-transferred i.v. into separate naïve recipients. Two weeks after transfer, the mice were infected i.v. with ≥2 × 10^6^ PFU (high dose) of LCMV Clone 13 and analyzed 7 days later. The LCMV Armstrong and Clone 13 strains were propagated on BHK-21 or Vero cells respectively, and virus stocks were titrated by standard immunofocus assays on MC57G cells. To assess viral titers, organ samples were titrated with the standard immunofocus assays on MC57G cells(15).

### 2.3 Stainings and flow cytometry

To exclude dead lymphocytes, cells were labeled with a LIVE/DEAD fixable dye (BioLegend). For intracellular transcription factor stainings (T-bet, Bcl-6, TCF1, TOX, c-Maf, Ki-67), cells were fixed and stained at 4°C using the FoxP3/Transcription Factor Staining Buffer Set (eBioscience) according to the manufacturer’s instructions. For cytokine detection (IFN-γ, TNF-α, IL-2), cells were restimulated with PMA (5ng/ml, Sigma-Aldrich) and ionomycin (5μg/ml, Sigma-Aldrich) or with endogenous antigen-presenting cells (APCs) loaded with GP64-80 peptide (1μg/ml) for 4 hours with addition of brefeldin A (5μg/ml, Sigma-Aldrich) after 30 min. To assess intracellular cytokines, cells were fixed with 2% formaldehyde (Merck) at RT after restimulation and stained in PBS/BSA containing 0.05% Saponin (Sigma-Aldrich). Antibodies were purchased from BD Biosciences, BioLegend, eBioscience, and Miltenyi Biotec or produced in-house at the DRFZ. For detailed information on antibodies see Supplementary Materials. To assess cell apoptosis, cells were stained with Annexin V and Propidium iodide (PI) using the Annexin V binding buffer (eBioscience). When indicated, frequencies or geometric mean (GM) of sorted cell subsets were normalized to those of their respective mock cells (after LCMV Armstrong infection) or average of mock cells from each experiment (after LCMV Clone 13 infection) for protein quantification.

### 2.4 Bulk RNA and ATAC sequencing

Total RNA isolation of *in vivo*-differentiated CD4^+^ T cells sorted according to T-bet ZsGreen expression (d10 p.i. with LCMV Armstrong) was performed using Nucleospin RNA XS Micro Kit. RNA quality (RQN >8) was assessed with a Fragment Analyzer (Advanced Analytical) and quantified with a high sensitivity dsDNA Qubit assay (Invitrogen) before the cDNA library was prepared with the SMART-Seq v4 Ultra Low Input RNA Kit (Clontech) and the Nextera XT DNA library prep. reference guide (Illumina). Paired-end sequencing (2×76nt) of the libraries was performed with a NextSeq2000 device (Illumina). The obtained reads were mapped to the mm39 genome (annotation release: M27_GRCm39) using Hisat2 (PMID: 25751142) with default settings and quality was controlled with Trimmomatic package (16,17). Read counts were determined with featureCounts (18). DESeq2 (PMID:25516281) in Rstudio (Version 1.4.1717) was used for differential gene expression analysis (19). A gene was considered as differentially expressed, when log2FC > 1.0 or log2FC < -1.0 and P adjusted to < 0.05. For data visualization the following packages were used in RStudio: AnnotationDbi(20), pheatmap (21), EnhancedVolcano (22), and ggplot2 (23). Gene set enrichment analysis (GSEA) was conducted using clusterProfiler (24). For DNA isolation and library preparation the ATAC Seq Kit (Active Motif) was used according to manufacturer’s instructions. The libraries were quantified using the KAPA Library Quantification Kit (Kapa Biosystems) and were paired-end sequenced (2×76nt) with a NextSeq 2000 device (Illumina). QC and analysis on ATAC-seq libraries was performed using AIAP pipeline (25). The reads were mapped to mouse mm10 genome. The generated peaks files for each library were annotated using Homer package (26). The normalized peaks were visualized using WashU Epigenome Browser (27).

### 2.5 Statistical analysis

Statistical analysis was performed in GraphPad Prism (v7 and v9.0.0). All samples were tested for normality with Shapiro Wilk or D’Agostino and Pearson tests depending on the sample size. If samples passed normality, paired (d10 p.i. LCMV Arm) or unpaired *t* test (d7 p.i. LCMV Cl13) was performed. If samples failed normality testing, Wilcoxon test (d10 p.i. LCMV Arm) or Mann Whitney U-test (d7 p.i. LCMV Cl13) were performed. Samples were always compared individually to T-bet^High^ samples. To assess correlation of two proteins, simple linear regression analysis was performed. Viral titers were log converted prior to analysis. P values ≥ 0.05 were considered non-significant (ns). P values < 0.05 were considered significant. *P < 0.05; **P < 0.01; ***P < 0.001; ****P < 0.0001.

### 2.6 Data availability

Raw and processed RNA and ATAC sequencing datasets generated in this study have been deposited in the gene expression omnibus (GEO) database under the accession number GSE199981.

## 3 Results

### 3.1 Antiviral T-bet^+^ CD4^+^ T cells exhibit phenotypic intrapopulation heterogeneity

To assess phenotypic differences in the antiviral T-bet^+^ CD4^+^ T cells pool in detail, we utilized T-bet ZsGreen reporter mice (13). Recipient mice (Thy1.2^+^, also T-bet ZsGreen^+^ to prevent rejection) that had received naïve LCMV-specific (Smarta) CD4^+^ T cells from T-bet ZsGreen donors (Thy1.1^+^) were infected with LCMV Armstrong (Arm, 200 PFU) to elicit an acute viral infection. On day 10 post infection (p.i.), the now differentiated progeny of the transferred T-bet ZsGreen Smarta CD4^+^ T cells were re-isolated. T-bet^High^ and T-bet^Low^ subpopulations were analyzed separately by electronic gating (egating) and compared to all T-bet ZsGreen reporter expressing Smarta cells (T-bet^Mock^) as reference population (Fig 1A). The quantity of T-bet protein was additionally measured by intracellular staining and compared to the quantity of ZsGreen expression (Fig 1B). The vast majority of the cells (>95%) had acquired an effector or effector memory phenotype (CD44^+^CD62L^-^). Notably, T-bet^Low^ cells had a minor, but significant increase in cells with a central memory phenotype (CD44^+^CD62L^+^) compared to the other populations (Suppl. Fig 1A). The expression of Th1-associated proteins, Ly6C, which has been shown to be important for the homing of T cells to lymphoid organs (28) and CXCR3, which is induced by T-bet and allows for homing to inflamed tissue (29), were further assessed. While all T-bet^+^ Smarta CD4^+^ T cells expressed CXCR3 to some extent as expected, the T-bet^High^ population included a CXCR3^Low^Ly6C^+^ subpopulation (Fig 1C). Ly6C expression correlated positively with T-bet expression levels as previously observed (30), causing the majority of T-bet^High^ cells to be Ly6C^+^, while half of the T-bet^Low^ cells were Ly6C^-^.

**Figure 1:**
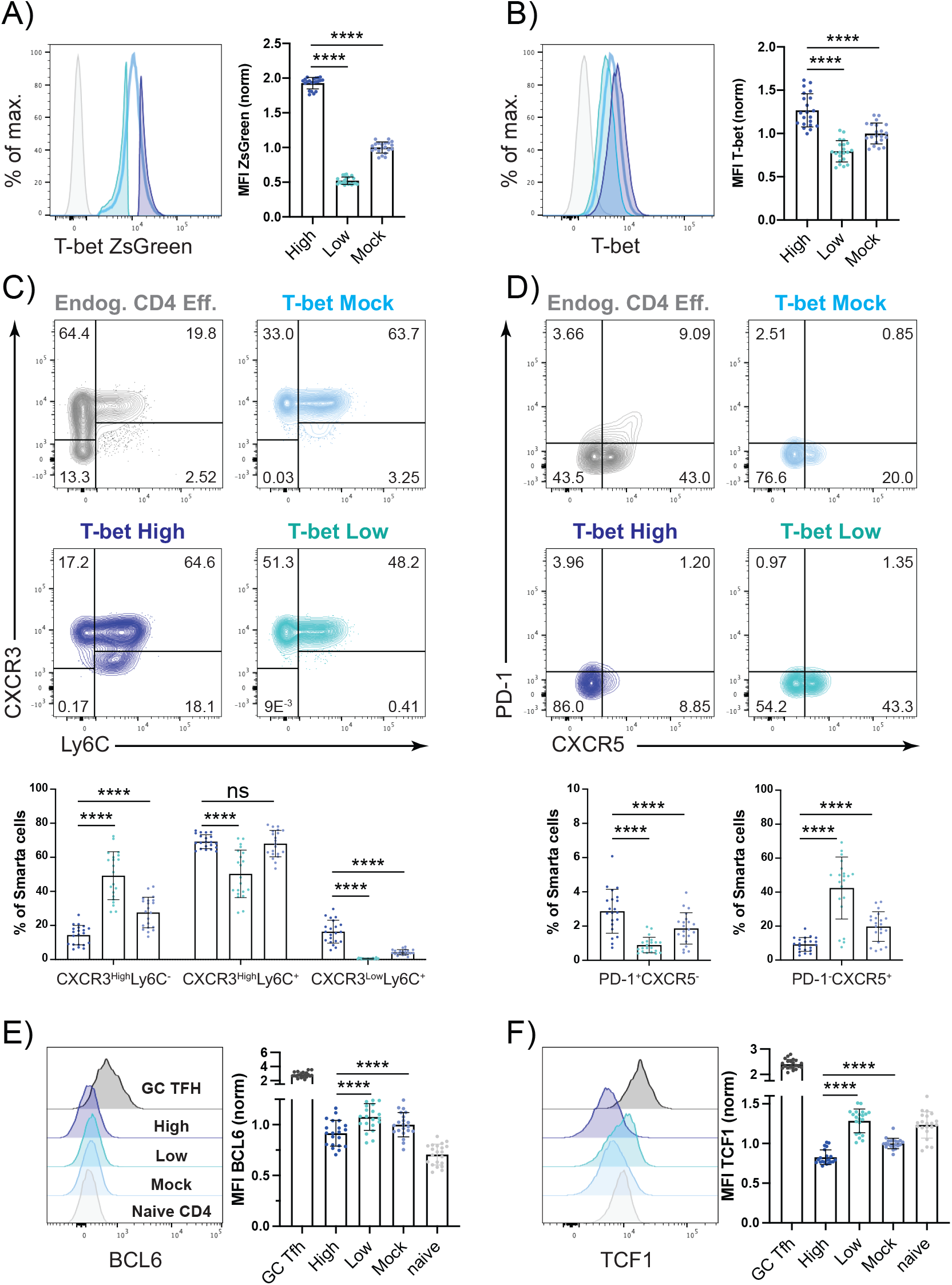
Antiviral T-bet^+^ CD4^+^ T cells exhibit phenotypic intrapopulation heterogeneity. Naive Smarta CD4^+^ T cells from T-bet ZsGreen donors (Thy1.1^+^) were transferred into T-bet ZsGreen recipients (Thy1.2^+^). Recipient mice were infected with LCMV Arm (200 PFU). On day 10 p.i. the cells were harvested from the spleen and lymph nodes. T-bet ZsGreen^+^ Smarta cells were electronically gated (egated) according to their T-bet reporter expression levels into T-bet^high^ or T-bet^low^ fractions, and all T-bet reporter positive cells (T-bet^mock^) served as controls. A) Representative histogram of T-bet ZsGreen expression (grey = naïve endog. CD4^+^ T cells). Pooled and normalized T-bet ZsGreen MFI of each egated fraction. B) Representative histogram of T-bet protein expression (grey = naïve endog. CD4^+^ T cells). Pooled and normalized T-bet protein MFI of each egated fraction. C) Representative gating of CXCR3 and Ly6C of endogenous effector CD4^+^ T cells (grey) or the egated T-bet^High^, T-bet^Low^ or T-bet^Mock^ fractions of Smarta cells. Pooled frequencies of different subsets. D) Representative gating of PD-1 and CXCR5 of endogenous effector CD4^+^ T cells (grey) or the egated fractions of Smarta cells. Pooled frequencies of different subsets. E) Representative histogram of BCL6 expression (grey = naïve endog. CD4^+^ T cells, dark grey = endogenous effector GC Tfh (PD-1^+^CXCR5^+^) cells). Pooled and normalized BCL6 MFI of each egated fraction. F) Representative histogram of TCF1 expression (grey = naïve endog. CD4^+^ T cells, dark grey = endogenous effector GC Tfh (PD-1^+^CXCR5^+^) cells). Pooled and normalized TCF1 MFI of each egated fraction. Data are presented as mean ± SD. Each dot represents isolated Smarta T cells or endogenous CD4^+^ T cells (grey) from one individual recipient. 5 independent experiments were pooled (n=4-5 mice/experiment). For MFI comparison, MFI of T-bet^High^, T-bet^Low^, endogenous GC Tfh or naïve CD4^+^ T cells were normalized to the corresponding T-bet^Mock^ sample. Statistical significance was determined by paired T-test or Wilcoxon test comparing T-bet low or mock to high Smarta cell fraction, statistical comparison to endogenous cells was not performed. p* <0.05, p**< 0.01, p****< 0.0001, ns = not significant.

During viral infections, also T follicular helper (Tfh) cells can arise, therefore we additionally assessed the expression of some Tfh-associated proteins in the T-bet positive virus-specific CD4^+^ T cells. We observed that up to half of the T-bet^Low^ compartment expressed low levels of CXCR5, a surface marker typically associated with Tfh cells and their homing to the B cell follicle (31) (Fig 1D). However, barely any CXCR5^+^ Smarta CD4^+^ T cells co-expressed PD-1, which has been shown to be important for Tfh cell positioning in the germinal center (GC) (32). The expression of the Tfh-associated transcription factors BCL6 and TCF1 was slightly, but significantly increased in T-bet^Low^ cells, yet remained at a much lower level than in endogenous GC Tfh cells (Fig 1E-F). These findings show that inside the T-bet^+^ virus-specific CD4^+^ T cells there is some phenotypic intrapopulation heterogeneity.

### 3.2 RNA-Seq identifies expression of various Tfh-associated genes preferentially in virus-specific Th cells with low T-bet expression

To further investigate potential differences between T-bet^High^ and T-bet^Low^ cells, we sorted the virus-specific cells according to their T-bet ZsGreen brightness on day 10 p.i. and performed RNA sequencing (Fig 2A). The cell populations showed a high overlap in gene expression with only up to 5% of the genes being significantly differentially expressed in cells with either high or low T-bet expression (Fig 2B). Analysis of the highly differentially expressed genes revealed a higher expression of some Tfh-associated genes (*Cxcr5, Tox2, P2r×7, Id3*) in T-bet^Low^ cells and cytotoxicity-associated genes (*Cx3cr1, Prf1, Gzmb*) in T-bet^High^ cells (Fig 2C). To further study the Tfh-related features at mRNA level in the T-bet^Low^ population, the gene set of Scholz et al. (33) for up- and downregulated genes in Tfh cells was used to perform enrichment scoring (Fig 2D). This indicated an enrichment of the Tfh signature upregulated genes in T-bet^Low^ and Tfh signature downregulated genes in T-bet^High^ cells (33). Furthermore, the expression of specific genes typically associated with either Th1 or Tfh cells was assessed (Fig 2E). T-bet^High^ cells showed higher expression of a number of Th1-associated genes, while T-bet^Low^ cells rather expressed higher levels of Tfh-associated genes. The majority of the Th1-associated genes was also highly expressed in T-bet^Low^ cells, emphasizing that even when there seems to be a bias towards some Tfh-like features, the T-bet^Low^ population can still be classified as part of the Th1 spectrum (Suppl. Table 1). This view is further supported by our observation of higher expression of *Ccr7* in T-bet^Low^ cells (Fig 2E), which is typically expressed by Th1 cells as it allows them to home to the T cell zone and has been shown to inhibit follicular homing of Tfh cells (34)(35). To assess whether some of the differences in gene expression could be explained by chromatin accessibility, DNA of the T-bet subgroups was analyzed by performing ATAC sequencing (Fig 2F). Overall, the chromatin accessibility showed only minor changes between T-bet^High^ and T-bet^Low^ samples (p-adj.<0.01: 573 out of 74980 unique peaks). Even though the *Tbx21* locus had similar accessibility in both subsets (data provided at GSE199981), we found changes at the transcription start sites of genes that were either higher expressed in T-bet^Low^ cells (*Cxcr5, Tox2*) or in T-bet^High^ cells (*Gzmb, Prf1*). Taken together, this points to some Tfh-like features in the antiviral T-bet^Low^ Th1 cell population.

**Figure 2:**
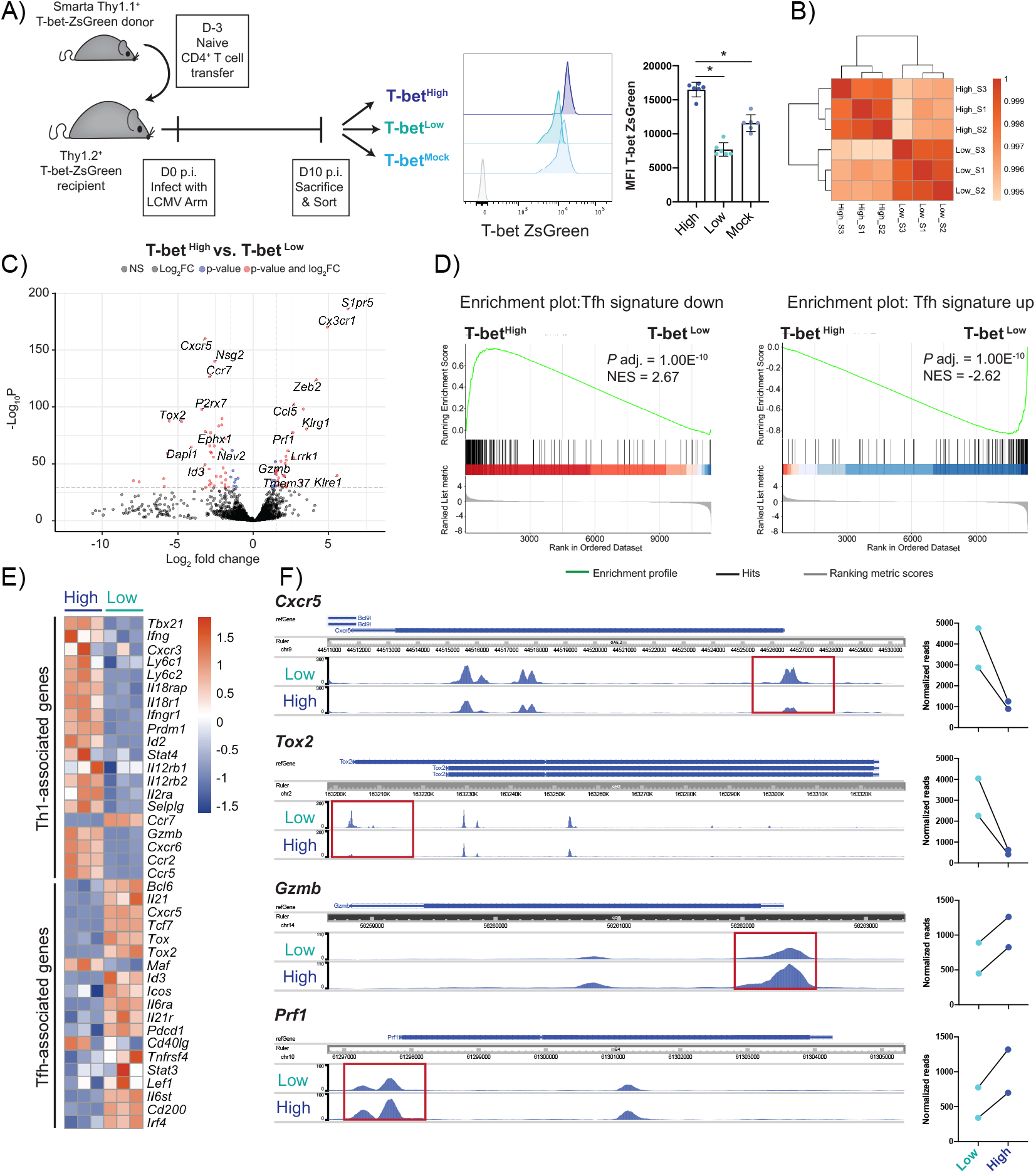
RNA-Seq identifies expression of various Tfh-associated genes preferentially in virus-specific Th cells with low T-bet expression. T-bet ZsGreen Smarta CD4^+^ T cells were differentiated as described in the previous figure. On day 10 p.i. with LCMV Arm, the cells were FACS sorted according to their T-bet reporter brightness into T-bet high, low and mock sorted fractions and bulk RNA- and ATAC-Sequencing was performed. A) Experimental outline, representative histogram of T-bet ZsGreen reporter MFI after sort (grey = naïve endogenous CD4^+^ T cells) and pooled T-bet ZsGreen MFI after sort of each fraction. B) Sample distance plot of shared and exclusively expressed genes in high vs. low T-bet ZsGreen CD4^+^ T cells. C) Volcano plot of T-bet reporter high vs. low sorted cell fractions. D) Gene set enrichment analysis of differentially regulated genes in the T-bet reporter sorted cells that have been shown to be either up- or downregulated in the Tfh cell signature (total 304 genes, gene set from publication by Scholz et al. 2021). E) Heatmap depicting the difference in expression of genes associated with either Th1 or Tfh phenotype in T-bet reporter high and low cells. F) ATAC Seq analysis of *Cxcr5, Tox2, Gzmb* and *Prf1* locus and normalized reads of highlighted significantly different peaks between T-bet reporter low and high cells. FACS data are presented as mean ± SD. Each dot represents T-bet ZsGreen sorted Smarta T cells from one experiment. 6 independent experiments were pooled (n=1/fraction/experiment). Samples of 2 (ATAC) or 3 (RNA) independent experiments were used for sequencing. Statistical significance was determined using paired T-test or Wilcoxon test comparing T-bet low or mock to high fraction. p* <0.05.

### 3.3 Quantitative differences in Th1 cell features are maintained to some extent during chronic viral rechallenge

To assess the stability of quantitative T-bet differences, adoptively transferred T-bet ZsGreen cells were sorted by reporter expression intensities on day 10 p.i. with LCMV Armstrong (Arm, 200 PFU), and the sorted cell fractions were transferred again into naïve Thy1.2^+^ recipients. After two weeks of resting, the recipients were challenged with LCMV Clone 13 (Cl13, ≥2 × 10^6^ PFU), which causes chronic infections in mice. As CD4^+^ T cells are usually recovered in low cell numbers during these infections, we decided to assess their phenotype in the spleen at an early time point (Fig 3A). Seven days p.i., the vast majority of transferred cells still expressed T-bet ZsGreen highly (Fig 3B). Even though the formerly T-bet ZsGreen brightness-sorted populations now showed an overlap in T-bet reporter expression, they still exhibited significant differences in their mean fluorescent intensity (MFI) (Fig 3B). To test the functional relevance of these differences, the levels of IFN-γ expression were assessed after *ex vivo* stimulation. While similar frequencies of the transferred cells expressed IFN-γ (Suppl Fig 2B), the T-bet^High^ cells expressed significantly more IFN-γ per cell than the T-bet^Low^ cells (Fig 3C). Furthermore, the originally observed differences in frequencies of CXCR3 and Ly6C expression patterns were maintained to some extent, as T-bet^Low^ cells still were mostly Ly6C^-^ (Fig 3D). However, the Ly6C expression of the transferred cells was overall decreased after LCMV Cl13 infection compared to the analysis after primary challenge with LCMV Arm. This observation was accompanied by slightly lower CXCR3 MFI and higher Ly6C MFI in the progeny of the T-bet^High^ compared to the T-bet^Low^ cells (Suppl Fig 2C). Additionally, IL-18R^+^ cell frequencies followed the T-bet ZsGreen gradient with highest percentages of IL-18R^+^ cells in the progeny of formerly T-bet^High^ sorted cells, further pointing to the potential functional relevance of the maintained quantitative T-bet ZsGreen differences (Suppl Fig 2D). IL-18 signaling has been shown to enhance IFN-γ production in differentiated Th1 cells (36). Thereby, heightened IL-18R expression may further strengthen the Th1 phenotype of T-bet^High^ cells. Taken together, quantitative differences in Th1 cells can be maintained to some extent even in a strong stimulatory and inflammatory environment as present during chronic viral infection.

**Figure 3:**
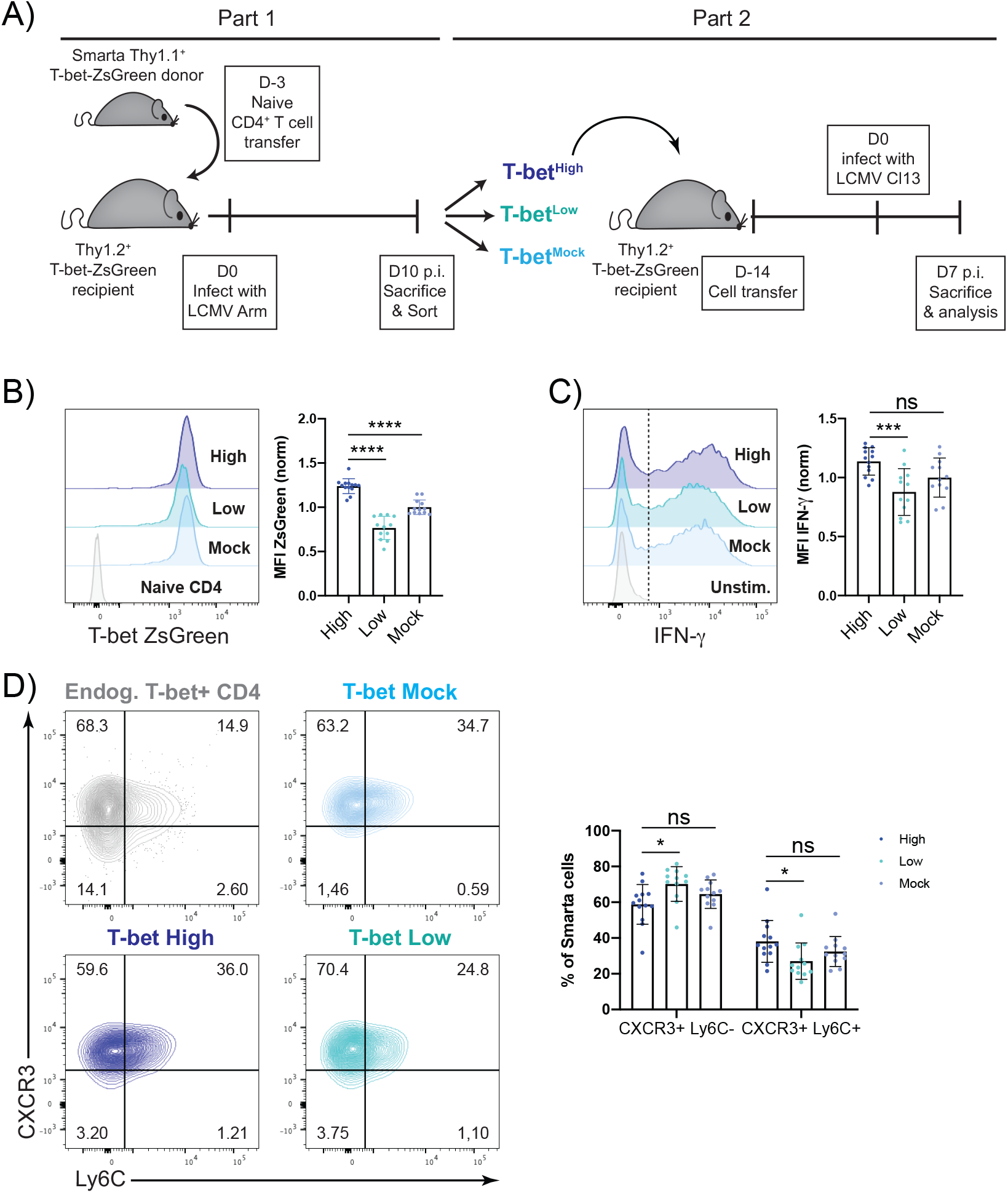
Quantitative differences in Th1 cell features are maintained to some extent during chronic viral rechallenge. Ten days p.i. with LCMV Arm, T-bet^High^, T-bet^Low^ and T-bet^Mock^ sorted Smarta cells (Thy1.1^+^) were transferred into individual naïve T-bet ZsGreen recipients (Thy1.2^+^). Two weeks post transfer, the recipients were infected with high dose LCMV Clone 13 (≥2×10^6^ PFU) and 7 days post infection, the transferred cells were isolated from spleen and their phenotype was analyzed with flow cytometry. A) Experimental Layout. B) Representative histogram of T-bet ZsGreen expression (grey = naïve endog. CD4^+^ T cells). Normalized and pooled T-bet ZsGreen MFI of transferred cells day 7 p.i. with LCMV Cl13. C) Representative histogram of IFN-γ expression after GP64 restimulation (grey = unstimulated T-bet^Mock^ cells). Normalized and pooled IFN-γ MFI of cytokine-positive transferred cells. D) Representative plots of CXCR3 and Ly6C staining of transferred cells and endogenous T-bet^+^ primary effector CD4^+^ T cells, as indicated. Pooled frequencies of CXCR3^+^ Ly6C^-^ and CXCR3^+^ Ly6^+^ Smarta cells. Data are presented as mean ± SD. Each dot represents isolated Smarta CD4^+^ T cells from one individual recipient. 3 independent experiments were pooled (n=4-5 mice/fraction/experiment). For MFI comparison, MFI of T-bet^High^ or T-bet^Low^ cells were normalized to the average of T-bet^Mock^ samples in each experiment. Statistical significance was determined using unpaired T-test or Mann-Whitney test comparing T-bet low or mock to high fraction. p* <0.05, p***< 0.001, p****< 0.0001, ns = not significant.

### 3.4 T-bet low Th1 cells preferentially maintain some Tfh-associated features and high T-bet expression does not prevent T cells from acquiring phenotypic markers of exhaustion

After the primary infection with LCMV Arm, the T-bet ZsGreen brightness-sorted cells showed significant differences in the expression of various classic Tfh-associated factors. Therefore, we wondered whether these features were maintained during chronic viral rechallenge. Already at day 7 p.i. with LCMV Cl13, the virus-specific secondary effector cells were all PD-1 positive (Fig 4A). The enrichment of CXCR5^+^ cells, which now co-expressed PD-1, was still slightly but significantly higher in the progeny of the T-bet^Low^ than the T-bet^High^ population. Even though the overall frequencies of CXCR5^+^ cells were reduced, the progeny of the transferred T-bet^Low^ cells still showed slightly but significantly higher expression of some Tfh-associated transcription factors including BCL6, TCF1, and c-Maf (Fig 4B). Overall, we could observe high expression of c-Maf in all subfractions confirming that this transcription factor is not only expressed in Tfh and Th2 cells (37–39) but also in chronically stimulated T cells as previously reported (40). The expression of the transcription factor TOX, which has been associated with the persistence of antiviral CD8^+^ T cells in chronic infections (41,42), was also assessed in the progeny of the T-bet reporter cells. After primary challenge, we found a mild increase in TOX expression in T-bet^Low^ cells (Suppl Fig 1C). However, after Cl13 challenge, we observed high expression of TOX, which was then comparable in the progeny of all transferred cell fractions (Fig 4B). Even though TOX and TOX2 have been shown to be involved in Tfh development, our data indicate that in a chronic infection setting, the transcription factor is not exclusive for Tfh cells, but rather strongly expressed by all antiviral CD4^+^ T cells (43).

**Figure 4:**
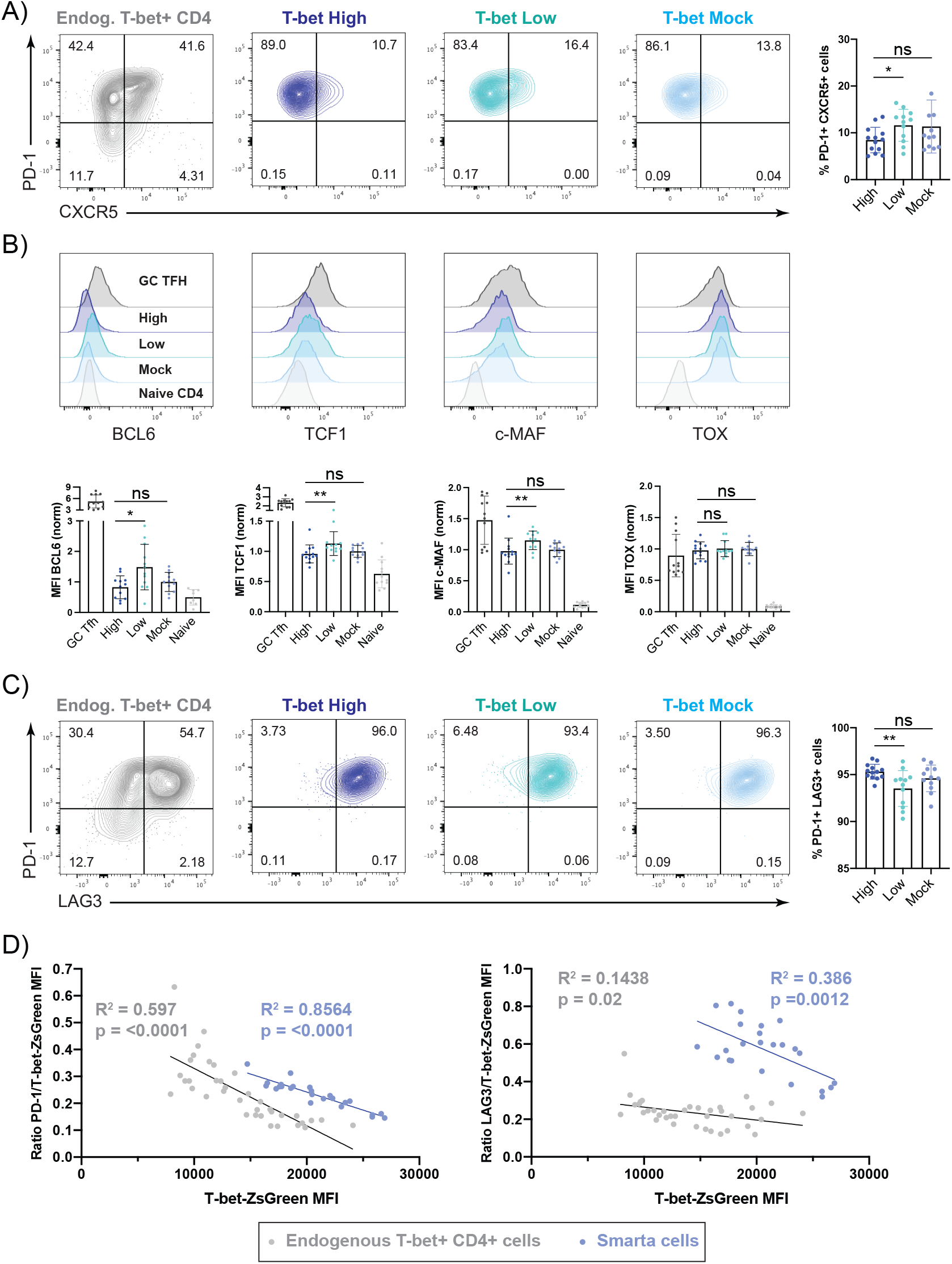
T-bet low Th1 cells preferentially maintain some Tfh-associated features and high T-bet expression does not prevent T cells from acquiring phenotypic markers of exhaustion. Experimental Layout as described in Figure 3. A) Representative plots of PD-1 and CXCR5 gating of Smarta cells (shades of blue) or endogenous T-bet^+^ effector CD4^+^ T cells (grey) in the spleen. Pooled frequency of PD-1^+^ CXCR5^+^ Smarta cells. B) Representative histograms of BCL6, TCF1, c-MAF and TOX expression levels in Smarta cells (shades of blue) in the spleen. Endogenous naïve CD4^+^ T cells (light grey) and endogenous effector GC Tfh (PD-1^+^CXCR5^+^) cells (dark grey) are depicted as controls. Normalized and pooled MFIs of BCL6, TCF1, c-MAF or TOX. C) Representative plots of PD-1 and LAG3 gating of Smarta cells or endogenous T-bet^+^ effector CD4^+^ T cells in the spleen, as indicated. Pooled frequency of PD-1^+^LAG3^+^ Smarta cells. D) Correlation analysis of PD-1 (left) or LAG3 (right) MFI ratio to T-bet ZsGreen MFI of either endogenous T-bet^+^ effector CD4^+^ T cells (grey) or transferred Thy1.1^+^ Smarta CD4^+^ T cells (blue). R2 and p value are stated in the graphs in the respective colors. Data are presented as mean ± SD. Each dot represents isolated Smarta CD4^+^ T cells from one individual recipient or endogenous CD4^+^ T cells from individual T-bet^mock^ cell recipients. 3 independent experiments (2 for Figure 4D) were pooled (n=4-5 mice/fraction/experiment). For MFI comparison, MFI of T-bet^High^, T-bet^Low^, endogenous GC Tfh or naïve CD4^+^ T cells were normalized to the average of T-bet^Mock^ samples in each experiment. Statistical significance was determined by unpaired T-test or Mann-Whitney test comparing T-bet Low or Mock to High fraction, statistical comparison to endogenous cells was not performed. p* <0.05, p**< 0.01, ns = not significant.

Next, we investigated whether the progeny of the sorted cell subsets with high or low T-bet reporter expression showed any differences in inhibitory receptor expression during chronic infection. As mentioned before, the secondary effector cells were all PD-1 positive, and LAG3, an inhibitory receptor competing with CD4 for binding, was expressed by almost all of the transferred cells (High: 95.3% +/-0.8, Low: 93.52% +/-1.91, Mock: 94.6% +/-1.41) (Fig 4C) (44,45). Here, T-bet^High^ cells showed a significant, but minor increase in PD-1^+^LAG3^+^ cell frequencies. As T-bet had been previously shown to be able to bind the *Pdcd1* locus and to repress PD-1 expression in CD8^+^ T cells (46)we assessed the PD-1 expression levels in activated CD4^+^ T cells after chronic LCMV infection. Even though the majority of T-bet ZsGreen^+^ endogenous effector and transferred secondary effector CD4^+^ T cells were PD-1^+^, the expression levels of the T-bet reporter negatively correlated with those of PD-1 in both cell types (Fig 4D). This suggests that T-bet can partially repress PD-1 in exhausted CD4^+^ T cells. However, as T-bet levels showed a weaker negative correlation with LAG3, T-bet does not seem to repress inhibitory receptor expression globally.

Furthermore, low proliferation rates, high cell death, and poor polyfunctionality are common features of T cell exhaustion (47,48), which we also observed in our transferred cells. The progeny of the sorted cells with different T-bet reporter expression levels showed comparable frequencies of proliferating, dead, or apoptotic cells on day 7 p.i. (Suppl Fig 3A-C). Additionally, the majority of the progeny of the transferred cells had already lost the capacity to produce TNF-α or IL-2 in addition to IFN-γ by day 7 p.i. showing no significant differences between the secondary effector cell populations (Suppl Fig 3D). Finally, as our analysis was done during the peak of the LCMV replication and as CD4^+^ T cells are not the main players in viral clearance (48), we also did not observe any differences in viral titers between the recipients (Suppl Fig 3E).

In summary, the progeny of the transferred CD4^+^ T cell fractions sorted according to T-bet reporter levels all exhibit multiple phenotypical features of exhaustion by day 7 of infection with LCMV Cl13. T-bet does not appear to globally prevent exhaustion in virus-specific Th1 cells, however higher T-bet reporter expression is correlated with lower PD-1 expression levels in individual cells.

## 4 Discussion

T-bet expression is considered a hallmark of CD4^+^ T cell differentiation towards the Th1 phenotype (49). However, in recent years it has become increasingly clear that not only the presence of a transcription factor, but its expression level in single cells can be decisive for cell-fate decisions and plasticity of differentiated cells (50,51). In the present study, we showed that within the T-bet-expressing antiviral CD4^+^ T cell compartment there is a continuous spectrum from T-bet^high^ to T-bet^low^ cells, which show intrapopulation heterogeneity. Furthermore, we could demonstrate that cells sorted according to the intensity of the T-bet ZsGreen reporter can maintain quantitative differences in T-bet reporter expression and IFN-γ expression even after a rechallenge with a chronic LCMV strain. Even though the cells maintained differential expression profiles, neither of the sorted populations were particularly protected from acquiring phenotypic markers of exhaustion during the early days of chronic infection.

We have previously reported that antiviral CD4^+^ T cells retain a quantitative memory of IFN-γ expression for at least one month after transfer into naïve recipients (1). Furthermore, we demonstrated that IFN-γ production is quantitatively controlled by T-bet, and these results were confirmed using the T-bet ZsGreen reporter (13). Moreover, we have recently demonstrated that the magnitude of T-bet expression safeguards Th1 plasticity (12). Here, we demonstrate that not only IFN-γ, but also homing markers (Ly6C, CXCR5), transcription factors (BCL-6, TCF1), and cytotoxic molecules (Perforin, Granzyme B) are influenced by the T-bet expression level in individual cells. Our data suggest that low T-bet amounts allow for increased expression of Tfh-associated markers. While a fraction of T-bet^low^ cells did express CXCR5, which allows T cells to enter the B cell follicle (52), they did not co-express PD-1 after the acute LCMV infection. PD-1 has been previously shown to be highly expressed by germinal center Tfh cells and to be able to restrict CXCR3 expression in CD4^+^ T cells (32,35). However, the T-bet^low^ cells strongly expressed CXCR3 and, according to the RNA-sequencing data, even upregulated *Ccr7* in comparison to T-bet^high^ cells, which has been shown to be able to inhibit follicular homing of CXCR5^+^ T cells (35). Even though Th1-like Tfh cells have been described before, these were rather defined by the early transient expression of T-bet, which resulted in the increased accessibility of the *Ifng* locus and thereby allowed for IFN-γ expression by the Tfh cells at later stages in the absence of T-bet (4). As the cells described here still expressed T-bet strongly after the clearance of the virus and only show a mild expression of BCL-6, we hypothesize that they are Th1 cells that in addition have certain Tfh characteristics. Previous studies in human blood have shown that PD-1^-^ CXCR3^+^ CXCR5^+^ CD4^+^ T cells, as we have observed in the T-bet^Low^ compartment, are IFN-γ producers, express T-bet, but are incapable of helping B cells (53). These cells were also described in the memory CD4^+^ T cell compartment of HIV patients and analysis of their expression profile neither matched the one from GC Tfh cells nor from CXCR5-negative T helper cells (54). It is possible that the T-bet^Low^ Th1 cells that feature certain Tfh characteristics described in the present study are the mouse equivalent, possibly located at the T:B border or, as the spleen is a highly vascularized organ, they might be circulating in peripheral blood. Further experiments will be required to define their exact localization.

The T-bet^high^ CD4^+^ T cells showed an upregulation of typical Th1 cell-associated genes and a strong expression and chromatin accessibility of the cytotoxic genes *Prf1* and *Gzmb*. It has been previously shown that T-bet can directly bind to the *Prf1* and the *Gzmb* genes in NK cells (55) and that the cytotoxicity of CD8^+^ T cells is impaired in T-bet KO strains (56). This in combination with our observations further suggests that high levels of T-bet may facilitate cytotoxic functions even in CD4^+^ T cells.

Even though we observed distinct protein and mRNA expression patterns in the T-bet reporter sorted subsets, these differences were not visible to the same extent in the ATAC-Seq data set. The global chromatin accessibility was comparable between samples, with only singular peaks differing significantly at the transcription start sites of highly differentially expressed genes as *Cxcr5, Tox2, Gzmb*, and *Prf1*. As the *Tbx21* and *Ifng* locus showed similar chromatin patterns (data provided at GSE199981), we hypothesize that the fine-tuned differences of their expression levels are rather mediated by transcription factor availability than by gene accessibility.

During chronic viral infections, as in tumors, T cells undergo functional adaptation that may include exhaustion due to the strong and continuous antigenic stimulation and the inflammatory environment (8). Several characteristics have been established to be part of the T cell exhaustion phenotype, some of which we observed in the progeny of all T-bet reporter sorted cell fractions after LCMV Cl 13 challenge infection: The cells became highly apoptotic, showed low proliferative potential and poor cytokine polyfunctionality (47,48,57). Furthermore, the progeny of the T-bet reporter-sorted CD4^+^ T cells showed high expression of the inhibitory receptors PD-1 and LAG3 already after 7 days post infection with LCMV Cl 13. While the vast majority of the transferred cells was positive for both markers, we still observed a strong negative correlation between the T-bet reporter expression and PD-

1 expression. This is in line with the findings by Kao et al., who had shown that in CD8^+^ T cells T-bet can negatively regulate the expression levels of inhibitory receptors, in particular PD-1, which is also the case for CD4^+^ T cells (46). Furthermore, the transcription factors c-Maf and TOX, which are associated with CD8^+^ T cell exhaustion, were strongly upregulated in the transferred T-bet reporter-sorted CD4^+^ T cell populations (40–42). Although both proteins have also been associated with Tfh cells, their strong upregulation during the chronic infection shows that they are not limited to the Tfh phenotype (37,38,43). Further studies are required to decipher the exact role particularly of TOX expression in exhausted CD4^+^ T cells during chronic LCMV infection.

In conclusion, our findings highlight the stability of graded quantitative levels of T-bet and IFN-γ expression in individual Th1 cells during chronic viral infection and the role of T-bet amounts in fine-tuning the expression of certain Tfh-associated genes in virus-specific Th1 cells.

## 8 Author Contributions

VP, CP, and ML designed the experiments. VP, AMA, ND, PS, VH, and IP performed the research. VP and MD analyzed the data. KL and MFM sequenced the samples. JZ provided T-bet ZsGreen reporter mice and advice. ANH provided advice on experimental design. VP, AMA, CP, and ML wrote the paper.

## 10 Acknowledgments

We thank C. Rüster for expert technical assistance, and L. Bauer for helpful discussions. Cell sorting was carried out at the Flow Cytometry Core Facility at the DRFZ.

## 11 Data Availability Statement

The datasets for raw and processed RNA and ATAC sequencing data generated for this study can be found in the gene expression omnibus (GEO) database under the accession number GSE199981. https://www.ncbi.nlm.nih.gov/geo/query/acc.cgi?acc=GSE199981 (publicly available before publication).

