## Supplementary material for "Stable characteristics of intrapopulation heterogeneity in virus-specific Th1 cells during chronic viral challenge infection"


#### 1 Supplementary Figures

1.1 Supplementary Figure 1: Phenotypic features of antiviral T-bet$^{+}$ primary effector T cells

1.2 Supplementary Figure 2: Th1 characteristics of the progeny of antiviral T-bet reporter-sorted CD4$^{+}$ T cells after LCMV Clone 13 challenge

1.3 Supplementary Figure 3: Phenotypic markers of exhaustion assessed in the progeny of antiviral T-bet reporter-sorted CD4$^{+}$ T cells after LCMV Clone 13 challenge

#### 2 Supplementary Methods

2.1 Antibody list

### 1 Supplementary Figures

#### 1.1 Supplementary Figure 1: Phenotypic features of antiviral T-bet$^+$ primary effector T cells

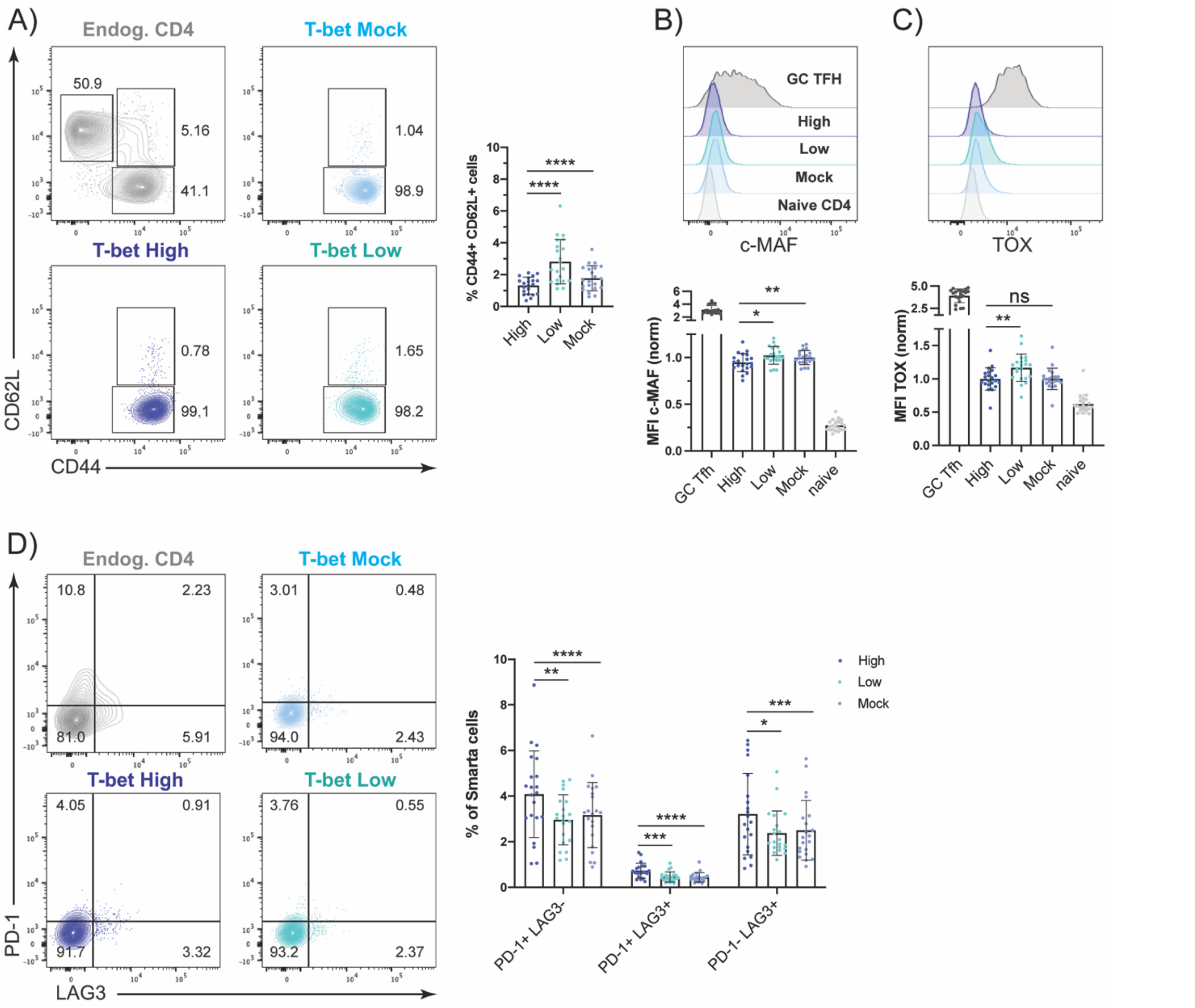


**Supplementary Figure 1.** Naïve Smarta CD4$^+$ T cells from T-bet ZsGreen donors (Thy1.1$^+$) were transferred into T-bet ZsGreen recipients (Thy1.2$^+$). Recipient mice were infected with LCMV Arm (200pfu). On day 10 p.i. the cells were harvested from the spleen and lymph nodes. T-bet ZsGreen$^+$ Smarta cells were electronically gated (egated) according to their T-bet reporter expression levels into T-bet$^{high}$ or T-bet$^{low}$ fractions, and all T-bet reporter positive cells (T-bet$^{mock}$) served as controls. (A) Representative gating of CD44 and CD62L of endogenous CD4$^+$ T cells (grey) or the T-bet reporter sorted Smarta cells (shades of blue). Pooled CD44$^+$CD62L$^+$ frequencies of Smarta cells. (B) Representative histogram of c-MAF expression (grey = naïve endog. CD4$^+$ T cells, dark grey = endogenous effector GC Tfh (PD-1$^+$CXCR5$^+$) cells). Pooled and normalized c-MAF MFI of each egated fraction. (C) Representative histogram of TOX (grey = naïve endog. CD4$^+$ T cells, dark grey = endogenous effector GC Tfh (PD-1$^+$CXCR5$^+$) cells). Pooled and normalized TOX MFI of each egated fraction. (D) Representative gating of PD-1 and LAG3 of endogenous CD4$^+$ T cells (grey) or the egated fractions of Smarta cells (shades of blue). Pooled frequencies of different subsets. Data are presented as mean ± SD. Each dot represents isolated Smarta T cells or endogenous CD4 T cells (grey) from one individual recipient. 5 independent experiments were pooled (n=4-5 mice/experiment). For MFI comparison, MFI of T-bet$^{High}$, T-bet$^{Low}$, endogenous GC Tfh or naïve CD4 T cells were normalized to the corresponding T-bet$^{Mock}$ sample. Statistical significance was determined by paired T-test or Wilcoxon test comparing T-bet low or mock to high Smarta cell fraction, statistical comparison to endogenous cells was not performed. p* <0.05, p**< 0.01, p***< 0.001, p****< 0.0001, ns = not significant.

#### 1.2 Supplementary Figure 2: Th1 characteristics of the progeny of antiviral T-bet reporter-sorted CD4$^+$ T cells after LCMV Clone 13 challenge

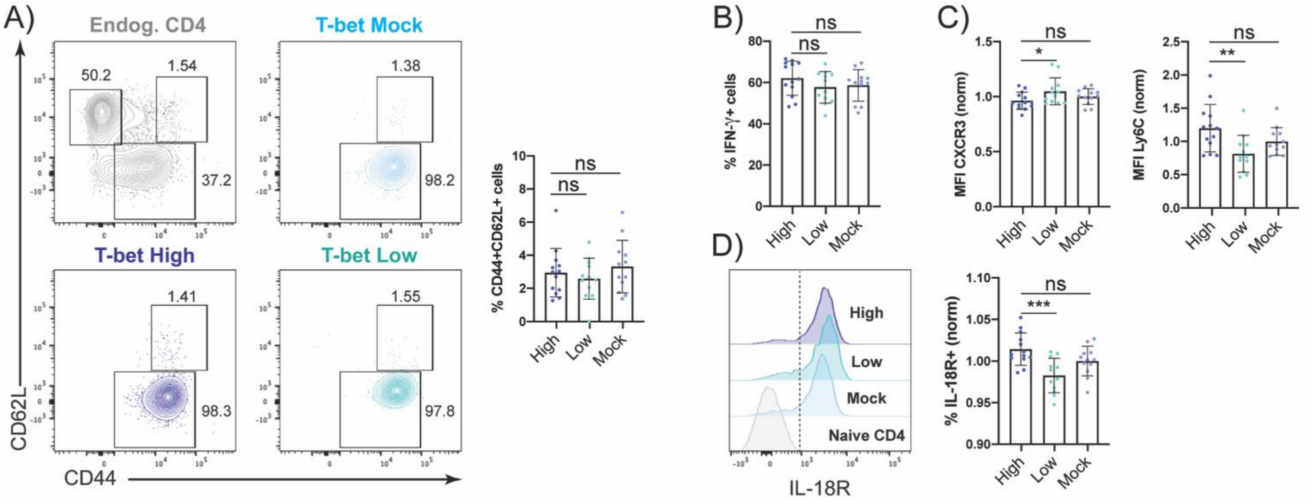


**Supplementary Figure 2.** Ten days p.i. with LCMV Arm, T-bet$^{High}$, T-bet$^{Low}$ and T-bet$^{Mock}$ sorted Smarta cells (Thy1.1$^+$) were transferred into individual naïve T-bet ZsGreen recipients (Thy1.2$^+$). Two weeks post transfer, the recipients were infected with high dose LCMV Clone 13 ($\geq 2 \times 10^6$ pfu) and 7 days post infection, the transferred cells were isolated from spleen and their phenotype was analyzed with flow cytometry. (A) Representative gating of CD44 and CD62L of endogenous CD4$^+$ T cells (grey) or Smarta cells (shades of blue). Pooled frequency of CD44$^+$CD62L$^+$ Smarta cells. (B) Pooled frequencies of IFN-γ$^+$ Smarta CD4$^+$ T cells after GP64-restimulation *ex vivo*. (C) Normalized and pooled MFI of CXCR3 and Ly6C of Smarta CD4$^+$ T cells. (D) Representative histogram of IL-18R expression of Smarta CD4$^+$ T cells (shades of blue) or naïve endogenous CD4$^+$ T cells (grey). Normalized and pooled frequencies of IL-18R$^+$ Smarta CD4$^+$ T cells. Data are presented as mean ± SD. Each dot represents isolated Smarta T cells from one individual recipient. 3 independent experiments were pooled (n=4-5 mice/fraction/experiment). For MFI or IL-18R$^+$ % comparison, MFI or frequencies of T-bet$^{High}$ or T-bet$^{Low}$ cells were normalized to the average of T-bet$^{Mock}$ samples in each experiment. Statistical significance was determined by unpaired T-test or Mann-Whitney test comparing T-bet low or mock to high fraction. $p^* < 0.05$, $p^{**} < 0.01$, $p^{***} < 0.001$, ns = not significant.

#### 1.3 Supplementary Figure 3: Phenotypic markers of exhaustion assessed in the progeny of antiviral T-bet reporter-sorted CD4+ T cells after LCMV Clone 13 challenge

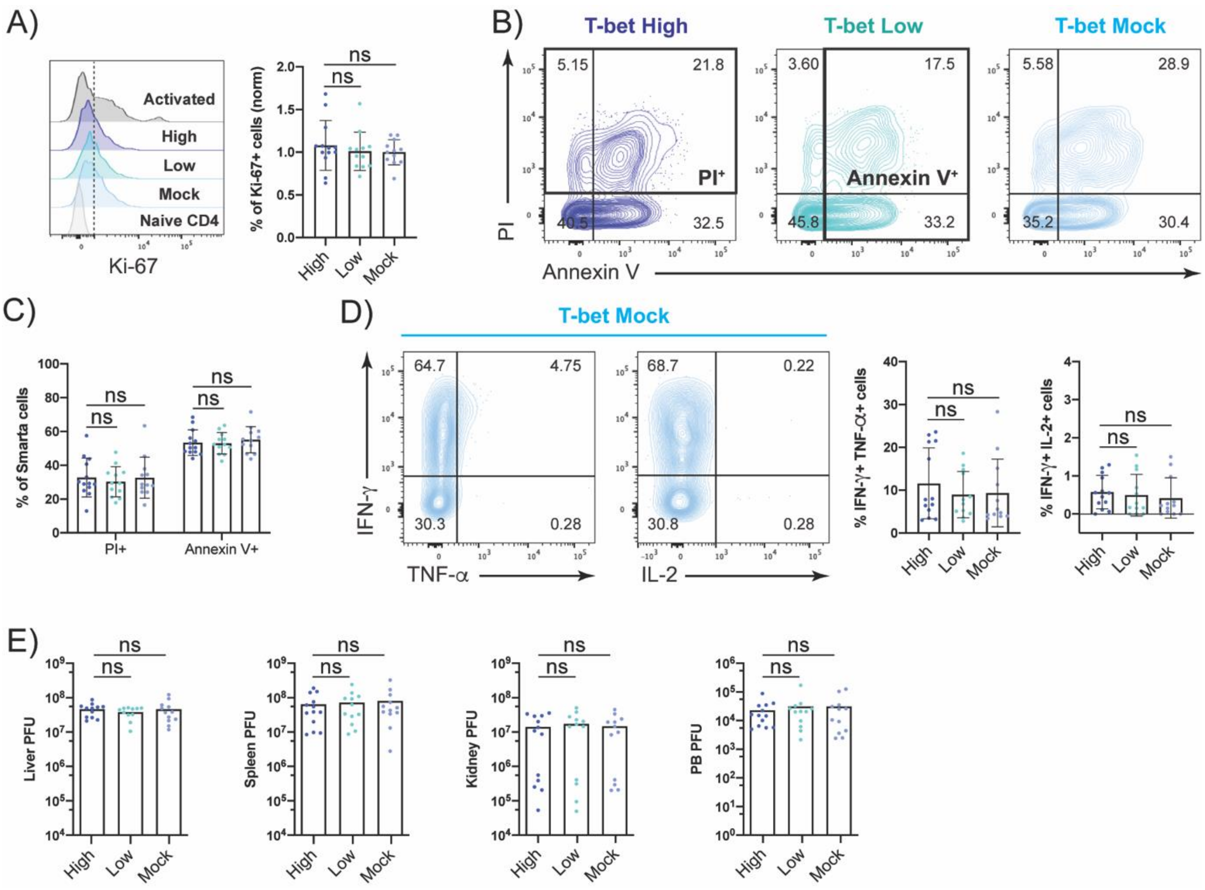


**Supplementary Figure 3.** Ten days p.i. with LCMV Arm, T-bet$^{High}$, T-bet$^{Low}$ and T-bet$^{Mock}$ sorted Smarta cells (Thy1.1$^{+}$) were transferred into individual naïve T-bet ZsGreen recipients (Thy1.2$^{+}$). Two weeks post transfer, the recipients were infected with high dose LCMV Clone 13 (≥2x10$^{6}$ pfu) and 7 days post infection, the transferred cells were isolated from spleen and their phenotype was analyzed with flow cytometry. (A) Representative histogram of Ki-67 expression of Smarta CD4$^{+}$ T cells (shades of blue) or endogenous naïve (light grey) or activated (dark grey) CD4$^{+}$ T cells. Normalized and pooled frequencies of Ki-67$^{+}$ Smarta CD4$^{+}$ T cells. (B) Representative gating of Annexin V and PI of Smarta CD4$^{+}$ T cells. (C) Pooled frequencies of dead (PI$^{+}$) or apoptotic (AnnexinV$^{+}$) Smarta cells. (D) Representative gating of IFN-γ and TNF-α or IL-2 coexpression of T-bet$^{Mock}$ Smarta CD4$^{+}$ T cells after GP64-restimulation *ex vivo*. Pooled frequencies of IFN-γ$^{+}$TNF-α$^{+}$ and IFN-γ$^{+}$IL-2$^{+}$ Smarta cells. (E) Pooled viral titers (plaque forming units, PFU) of liver, spleen, kidney and peripheral blood (PB). Data are presented as mean ± SD. Each dot represents isolated Smarta CD4$^{+}$ T cells from one individual recipient. 3 independent experiments were pooled (n=4-5 mice/fraction/experiment). For comparison of Ki-67$^{+}$ cell frequencies, frequencies of T-bet$^{High}$ or T-bet$^{Low}$ cells were normalized to the average of T-bet$^{Mock}$ samples in each experiment. Statistical significance was determined by unpaired T-test or Mann-Whitney test comparing T-bet low or mock to high fraction. ns = not significant.

### 2 Supplementary Methods

#### 2.1 Antibody list

| Target | Conjugate | Clone | Company | Cat. Number | RRID |
|---|---|---|---|---|---|
| Bcl6 | A647 | K112-91 | BD | 561525 | AB_10898007 |
| IL-18Ra | A647 | A17071D | BioLegend | 157907 | AB_2860736 |
| Ki-67 | A647 | B56 | BD | 561126 | AB_10611874 |
| T-bet | AF488 | 4B10 | Biolegend | 644829 | AB_2566018 |
| Thy1.1 | AF700 | OX-7 | Biolegend | 202527 | AB_1626244 |
| Annexin V | APC | - | BD | 550475 | AB_2868885 |
| LAG3 | APC | C9B7W | eBioscience | 17-2231-80 | AB_2573183 |
| PD-1 | APC | J34 | eBioscience | 17-9985-80 | AB_11149860 |
| TNF | APC | MP6-XT22 | BD | 561062 | AB_398553 |
| Thy1.2 | APC-Cy5 | Ho13 | In House | | |
| CD4 | APC-Cy7 | GK1.5 | BD | 565650 | AB_2739324 |
| CD44 | APC-Cy7 | IM7 | BioLegend | 103027 | AB_830784 |
| CXCR5 | Biotin | L138D7 | BioLegend | 145509 | AB_2562125 |
| CD8 | Biotin | 53-6.7 | In House | | |
| CD11b | Biotin | M1/70 | BD | 553309 | AB_394773 |
| CD11c | Biotin | HL3 | BD | 553800 | AB_395059 |
| CD25 | Biotin | 7D4 | In House | | |
| Gr-1 | Biotin | RB6-8C5 | BD | 553125 | AB_394641 |
| CD19 | Biotin | 1D3 | In House | | |
| CXCR3 | Biotin | CXCR3-173 | eBioscience | 13-1831-82 | AB_1210592 |
| NK1.1 | Biotin | PK136 | BD | 553163 | AB_394675 |
| T-bet | BV421 | 4B10 | Biolegend | 644832 | AB_2686976 |
| Tcf1/Tcf7 | BV421 | S22-966 | BD | 566692 | AB_2869822 |
| CD62L | BV605 | Mel-14 | Biolegend | 104437 | AB_11125577 |
| CD4 | BV650 | RM4-5 | BioLegend | 100545 | AB_11126142 |
| PD-1 | BV785 | 29F.1A12 | Biolegend | 135225 | AB_2563680 |
| IFNg | eFluor450 | XMG1.2 | eBioscience | 48-7311-80 | AB_1834367 |
| CXCR3 | PB | CXCR3-173 | Biolegend | 126529 | AB_2563100 |
| Thy1.1 | PB | OX-7 | In House | | |
| CD4 | PE | GK1.5 | BD | 553730 | AB_396634 |
| LAG3 | PE | C9B7W | Biolegend | 125207 | AB_2133344 |
| Tox | PE | TXRX10 | eBioscience | 12-6502-82 | AB_10855034 |
| PD-1 | PE | J43 | eBioscience | 12-9985-81 | AB_466294 |

| | | | | | |
|---|---|---|---|---|---|
| CD4 | PE-Cy7 | RM4-5 | BD | 552775 | AB_394461 |
| CD62L | PE-Cy7 | MEL-14 | eBioscience | 25-0621-82 | AB_469633 |
| IL2 | PE-Cy7 | JES6-5H4 | eBioscience | 25-7021-82 | AB_1235004 |
| Ly6C | PE-Cy7 | HK1.4 | eBioscience | 25-5932-82 | AB_2573503 |
| CXCR5 | PE-Vio770 | REA215 | Miltenyi | 130-117-366 | AB_2733206 |
| c-Maf | PerCP | sym0F1 | eBioscience | 46-9855-42 | AB_2573908 |
| CXCR3 | PerCP | CXCR3-173 | eBioscience | 45-1831-82 | AB_1210699 |
| Ly6C | PerCP | HK1.4 | eBioscience | 45-5932-82 | AB_2723343 |
| Streptavidin | PerCP | - | BD | 554064 | AB_2336918 |
| Thy1.1 (OX-7) | PerCP | OX-7 | BD | 557266 | AB_396611 |
| CD8 | V500 | 53-6.7 | BD | 560776 | AB_1937317 |
